# A novel mouse model of postpartum depression and the neurobiological effects of fast-acting antidepressant treatments

**DOI:** 10.1101/2022.05.06.490916

**Authors:** Alba García-Baos, Irene Ferreres-Álvarez, Inés Gallego-Landin, Xavier Puig-Reyné, Adriana Castro-Zavala, Olga Valverde, Ana Martín-Sánchez

**Affiliations:** Neurobiology of Behaviour Research Group (GReNeC-NeuroBio), Department of Medicine and Life Sciences, Universitat Pompeu Fabra, Barcelona, Spain; Neuroscience Research Program, IMIM-Hospital Del Mar Research Institute, Barcelona, Spain

**Keywords:** Allopregnanolone, GABA, glutamate, ketamine, maternal separation with early weaning, postpartum depression

## Abstract

postpartum depression (PPD) is a severe psychiatric disorder that affects up to 15% of mothers and impairs mother-infant bonding with devastating consequences on the child development and the mother health. Several studies indicate a possible dysregulation of glutamatergic and GABAergic signalling in the corticolimbic system, as well as a downregulation of the allopregnanolone levels in serum of PPD patients. Although brexanolone, an allopregnanolone-based treatment, has recently emerged as fundamental PPD treatment, there is scarce evidence on its neurobiological action mechanism. Moreover, ketamine appears to be a promising antidepressant treatment preventing PPD, nevertheless whether it might be a more effective than allopregnanolone for some patients remain unknown. Therefore, the present study is aimed to evaluate the depressive-like phenotype of postpartum females undergoing maternal separation with early weaning (MSEW) protocol, as well as to compare the effectiveness of ketamine and allopregnanolone treatments. MSEW dams show increased despair-like behaviour, anhedonia and disrupted maternal behaviour. Moreover, lower allopregnanolone serum levels, reduction of vesicular transporters for GABA (VGAT) and glutamate (VGLUT1) in the infralimbic cortex, as well as decreased hippocampal cellular proliferation are found in MSEW females. As for the antidepressant treatments, both drugs prevent despair-like behaviour, whereas only ketamine reverts anhedonia present in MSEW females. In addition, both treatments induce pro-neurogenic effects in the dorsal hippocampus but only allopregnanolone increases the VGAT and VGLUT1, without altering the excitatory/inhibitory ratio. Altogether, we propose a new mice model that recapitulates the core symptomatology and alterations in glutamatergic and GABAergic systems shown in PPD patients, which allows us to investigate the therapeutic mechanisms of allopregnanolone and ketamine.

## 1. Introduction

postpartum depression (PPD) is a subtype of Major Depressive Disorder (MDD) with peripartum onset which often appears within the first trimester after birth according to the Diagnostic and Statistical Manual of Mental Disorders (DSM-5) ^1^. This devastating psychiatric condition is characterised by feelings of worthlessness, insomnia, significant anxiety or maternal neglect ^2^. The prevalence of PPD is currently around 10-15% among women who given birth ^3,4^. However, this percentage is considered to be a misrepresentation since social stigma and other determinants might contribute to an overall underdiagnosis. In fact, the number of mothers reporting depressive symptoms rises up to 40% ^5,6^. Given such a large social burden, there is an imperative need to shed light on maternal mental health and the mother-infant bonding to address the underlying neurobiological mechanisms of PPD.

A decreased activation pattern in prefrontal cortex (PFC) and hippocampus (HPC), as well as other corticolimbic areas of PPD patients has been shown by functional magnetic resonance imaging studies ^7,8^. In fact, the reduction in neural connectivity observed in PPD ^9^ is probably due to an impairment in glutamatergic and GABAergic synapses, albeit they could be somehow balanced due to allostatic regulation ^9^. These results are supported by preclinical studies which demonstrate that chronic stress-based models of depression downregulate synaptic markers, like vesicular glutamate transporter (VGLUT) or vesicular GABA transporter (VGAT) among others, causing atrophy of these synapses and altering the neurotransmission within PFC and other corticolimbic regions ^10^. In addition to this, impairments of adult hippocampal neurogenesis might be a putative mechanism underlying functional hippocampal alterations shown in those patients. Preclinical models of depression exhibit impaired adult hippocampal neurogenesis ^11^, which has been associated with depressive-like behaviour observed in mice ^12^.

However, sex steroids affect many aspects of neuronal function that may contribute to the depression vulnerability. Fluctuations of reproductive-associated steroids, like oestrogen and progesterone, are known to increase the risk of depression ^13,14^. Therefore, it is essential to pay attention to critical periods of the reproductive life cycle, since the phases of high hormone fluctuations, such as pregnancy, overlap with at greater risk of developing depression ^15^. Thus, one might hypothesize that the molecular mechanisms underlying depressive symptoms during the peripartum period may be different compared to other types of major depression disorder. In particular, fluctuations of allopregnanolone have drawn particular in recent years. Allopregnanolone is an endogenous progesterone-derived neurosteroid synthetized in the brain that acts as a positive allosteric modulator of γ-aminobutyric acid ligand gated channel A (GABA_A_) receptors ^16^. In physiological conditions, allopregnanolone levels rise throughout pregnancy and peak around labour before rapidly returning to basal levels after birth ^9^. Nevertheless, PPD patients show low serum content of allopregnanolone in late pregnancy ^17^, leading to the hypothesis that reduced GABAergic signalling may underlie PPD symptomatology. Indeed, the hypofunction of GABA_A_ has been suggested to cause PPD-like behaviour in a mice model^18^.

Altogether the above-mentioned data supports the use of brexanolone, which is chemically identical to endogenous allopregnanolone and acts also as a positive allosteric modulator of GABA_A_ receptor, as a specific treatment for PPD ^19,20^. In truth, it is the only antidepressant drug employed for PPD that has been approved by the Food and Drug Administration ^21^.

Alternatively, ketamine has also been suggested as promising drug for preventing PPD in women who undergo caesarian sections ^22^. Ketamine is a glutamate N-methyl-d-aspartate (NMDA) non-competitive receptor antagonist that shows rapid antidepressant effects ^23^.

Thus, the approval of ketamine for treatment-resistant depressive patients ^24,25^ supports the idea of a promising effect on PPD as well.

Even though brexanolone and ketamine have both been approved and put forward respectively, the mechanisms through which they exert their beneficial effects on PPD patients remain unknown. In addition, the limited number of reliable PPD-like models that entirely mimic this complex psychiatric condition ^26^, may explain the misinformation referring to both the neuropathological basis of PPD and the antidepressants’ mechanism of action. For this reason, we have characterised a mouse model of PPD-like behaviour using the maternal separation with early weaning (MSEW). This model emulates human conditions better than other physical or pharmacological stressors since it employs both psychological and social stress when separating dams from their pups. Thus, the present study aimed to investigate whether MSEW dams show depressive-like symptomatology linked to correlated molecular markers: i) allopregnanolone plasma levels; ii) BDNF-TrkB pathway in PFC and HPC; iii) VGLUT1 and VGAT levels in infralimbic cortex (IL) as markers of glutamatergic and GABAergic neurotransmission; iv) adult hippocampal neurogenesis. Thereafter, we aimed to assess the effect of the two novel and most promising antidepressant drugs, ketamine and brexanolone, on PPD-like phenotype. Finally, to better understand the mechanisms of action underlying its effectiveness, we explore whether these treatments might restore the altered molecular correlates observed in the postpartum mothers undergoing MSEW model: i) GABAergic and glutamatergic signalling in IL, and ii) adult hippocampal neurogenesis.

## 2. Methods

### 2.1. Animals and rearing conditions

Male and female C57BL/6 mice were purchased from Envigo (Barcelona, Spain) to be used as breeders. Animals were housed with water and food *ad libitum* in a room maintained at 21 ± 1 °C, humidity 55 ± 10% and a12:12 h light-dark cycle, with lights off at 07:30 h, hence, behavioural experiments were performed during the dark phase under a dim red light. Breeding began when mice became 10 weeks old. For this, each male was housed together with 2 nulliparous females. After successful mating, pregnant females were individualized and observed daily for parturition.

For each litter, the date of delivery was designated as postpartum day (PD) 0. Litters of 6 pups were selected. Dams were randomly assigned to one of two rearing conditions: standard nest (SN) and maternal separation with early weaning (MSEW), which was conducted as previously ^27,28^. Briefly, MSEW litters were separated from their mothers for 4 h daily between PD 2-5 (09:30–13:30 h) and 8 h daily between PD6-16 (09:30–17:30 h), and then, they were weaned early on PD 17. For separation, mothers were removed to another cage while the offspring were maintained in their home cages with a heating blanket (32–34 °C) for thermoregulation. The SN offspring remained undisturbed with their mothers until the standard weaning day 21 (PD 21). After weaning, offspring were housed in groups of 4–5 animals of the same sex for another experiment and only dams were used for this study. All animal care and experimental procedures were conducted in accordance with the European Union Directive 2010/63/EU regulating animal research and were approved by the local Animal Ethics Committee (CEEA-PRBB).

### 2.2. Drugs

Ketamine hydrochloride (30 mg/kg; Imalgène1000, Lyon, France) and Allopregnanolone (Allop, 2mg/kg; Tocris, United Kingdom) were respectively diluted in a vehicle solution, composed by a mixture of (2-hydroxypropyl)-β- cyclodextrin (5%; Sigma Aldrich) and saline (0.9% NaCl). Both treatments were administered intraperitoneally (i.p.) in a volume of 0.1 mL/10 g of body weight. The doses of each treatment were chosen based on previous studies showing antidepressant effects ^29,30^.

### 2.3. Experimental design

For the characterization of PPD model (Experiment 1), forty-two animals were randomly assigned to either MSEW or SN experimental groups. In each rearing basal condition, twenty-four (SN n = 12, MSEW n = 12) postpartum females were subjected to the pup retrieval and dispersal test, the tail suspension test (TST) and the splash test to determine the maternal care, as well as the basal level of depressive-like and anhedonia-like behaviour. The rest of postpartum females were assigned to molecular studies (SN n = 9; MSEW n = 9; **Fig. 1A**).

**Fig.1.**
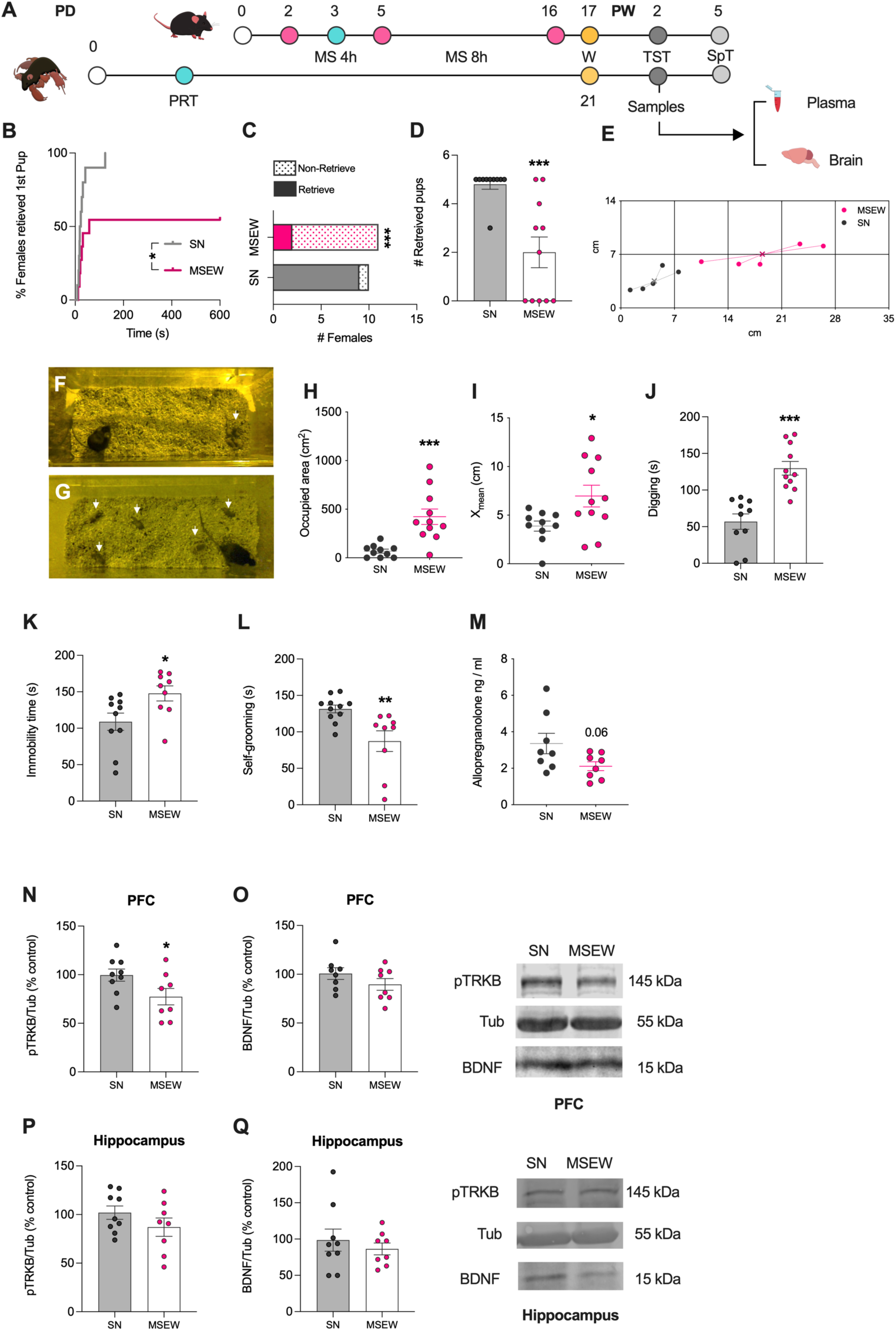
Characterization of MSEW as PPD mouse model. (A) Schematic representation of the experimental timeline. Grey (SN) and pink (MSEW) lines represent (B) the percentage of females that retrieved the first pup during the test. (C) Proportion of females that retrieved (filled bar) all pups. (D) Number of pups that were retrieved. Representative photographs of nests built by SN (F) and MSEW (G) mothers. (E) Dispersion of the pups by their respective dams is shown as the occupied area of the nest (H) and the average distance of the pups from the centroid from each litter (X_mean_; I) of SN and MSEW groups. (J) Time spent digging the sawdust. (K) Immobility time spent in TST and (L) self-grooming behaviour on splash test. (M) Content of allopregnanolone in plasma samples of SN and MSEW dams. pTRKB and BDNF protein levels in PFC (N,O) and hippocampus (P,Q). All data are represented as the mean ± SEM. (Log-rank test ^*^*p* < 0.05; Fischer’s test ^***^*p* < 0.001; t-Student’s test, ^*^*p* < 0.05, ^***^*p* < 0.001, Mann-Whitney’s test ^***^*p* < 0.001). MS, maternal separation; MSEW, maternal separation with early weaning; PD, postpartum day; PFC, prefrontal cortex; PRT, pup retrieval test; PW, postweaning day; SN, standard nest; SpT, splash test; TST, tail suspension test; W, day of weaning day.

In Experiment 2, dams were administered with ketamine (30 mg/kg, i.p.) 24 hours or allopregnanolone (2mg/kg, i.p.) 30minutes before each behavioural test and immunohistochemical analyses (**Fig.2A**). To counterbalance the injection timing, one half of the vehicle group received the injection either 24h or 30minutes prior the experiment. All treatments were randomly assigned to either SN or MSEW groups and were maintained for the entire battery of behavioural tests. The TST was carried out on postweaning day (PW) 2. On PW 5, splash test was performed. Following the behavioural evaluations, some animals (n=5-6/group) were randomly assigned to receive an injection of 0.9% NaCl v/v 5-bromo-2’Deoxyuridine (BrdU) 100 mg/kg (MERCK; New York, USA) three times within the same day, 24 hours or 30 minutes after receiving the last ketamine, allopregnanolone or vehicle injection, respectively. Finally, animals were perfused to further analysis of cellular proliferation in HPC, as well as glutamatergic and GABAergic signalling in IL on PW 12. The experimenters were blinded to both treatment and group while recording and analysing the behavioural procedures.

**Fig. 2.**
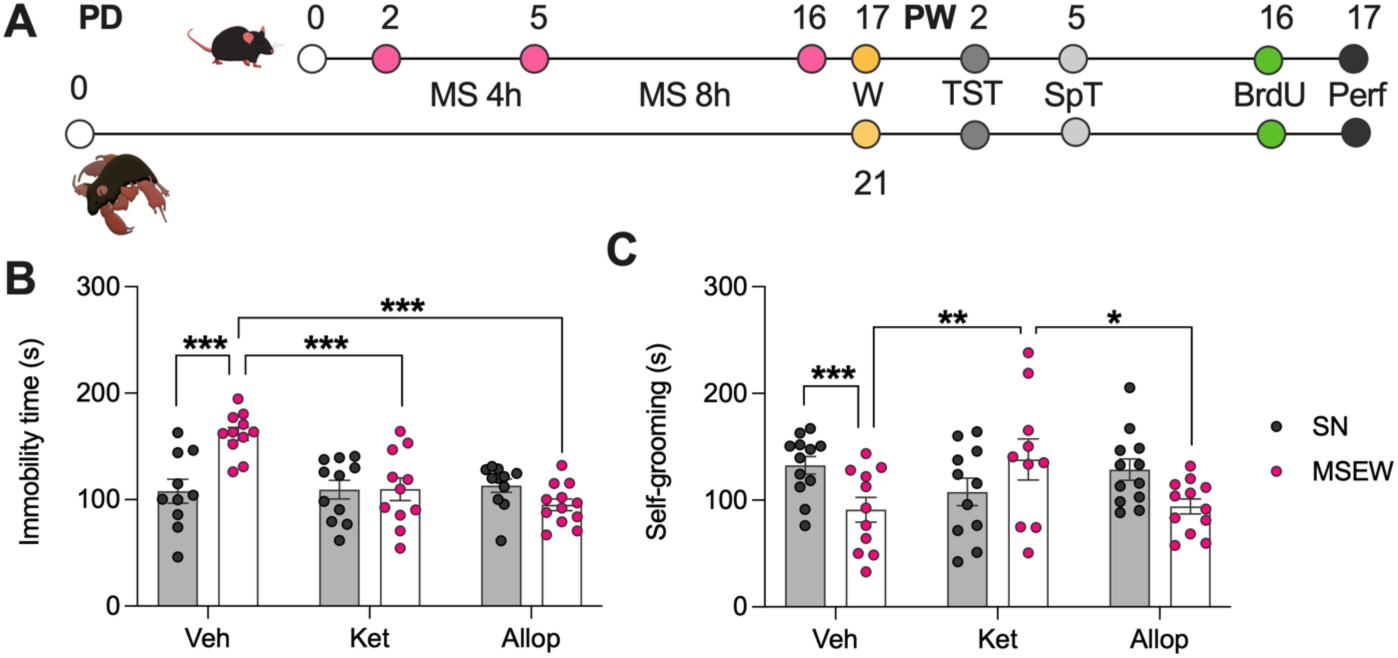
Differences between ketamine and allopregnanolone in reducing despair-like and anhedonia-like behaviours in MSEW females. (A) Timeline of the experiments. Grey (SN) and pink (MSEW) dots represent (B) the time that animals are immobile in TST after vehicle, ketamine or allopregnanolone treatments (n = 10-12 / group). (C) Graph shows the cumulative time self-grooming behaviour of SN (grey dots) and MSEW (pink dots) dams in splash test after vehicle, ketamine or allopregnanolone treatments (n = 10-12 / group). All data are represented as the mean ± SEM. (Two-way ANOVA ^*^p < 0.05, ^**^p < 0.01, ^***^p < 0.001). Allop, allopregnanolone; BrdU, 5-bromo-2’Deoxyuridine; Ket, ketamine; MS, maternal separation; MSEW, maternal separation with early weaning; perf, perfusion; PD, post-partum day; PRT, pup retrieval test; PW, post-weaning day; SN, standard nest; SPT, splash test; TST, tail suspension test; W, day of weaning day.

### 2.4. Pup retrieval and pup dispersal test

On PD 3, before MSEW dams were relocated into the home cage with their litters (n=10), five pups were scattered along the whole perimeter of the cage using a clean spatula as previously described ^31^. For SN females (n=10), the mother was removed from the home cage briefly (about 10 s), pups were dispersed as previously detailed. Then, the dam was reintroduced into the home cage. Every test was video-recorded for following analysis. We registered the latency to retrieve the first pup during the 600 s of the test and total time of digging the sawdust. If a female never retrieved it, a latency value of 600 s was assigned. Subsequently, the location of each pup was identified at minute 10 of the recording and it was evaluated by calculating the x and y coordinates, as reported ^18^.The centroid (mean of all x and all y values) was determined for each litter at this time point. The mean distance from the centroid for each pup (X_mean_) was calculated for every litter of SN and MSEW groups. Finally, the occupied area of each litter was calculated by using the ellipse area formula, Area = π · r_major_ · r_minor_, in which r_major_ and r_minor_ are the major and minor radius of the ellipse, respectively. The smallest distance from a pup to the centroid of its litter was designated as r_minor_, and the largest distance as r_major_.

### 2.5. Tail suspension test

On PW 2, each mouse was individually suspended from the tip of the tail, 50 cm above a bench top for 6 minutes. This test allows measuring total immobility time, reflecting despair-like behaviour ^32^ which in turn can be indicative of a depressive-like behaviour.

### 2.6. Splash test

Diminished self-grooming behaviour is an indicator of anhedonia in mouse models of depressive-like behaviour ^33^. On PW 5, the splash test was carried out in the female’s home cage to avoid stress of a new environment. Mice were placed in a corner of the home cage and sprayed on the back twice (∼2 × 0.6 ml) with 10% sucrose solution diluted in tap water ^33^. Total time of self-grooming behaviour (licking, stroking and scratching) were manually recorded for 5 minutes.

### 2.7. Western blot assay

postpartum females for each group were used for Western blot analysis and enzyme-linked immunosorbent assay (ELISA) (Experiment 1, **Fig. 1A**) on PWD2. Animals were anesthetised using a mixture of ketamine hydrochloride (75mg/kg; Imalgène1000, Lyon, France) and medetomidine hydrochloride (1mg/kg; Medeson ®, Barcelona, Spain) in a volume of 0.15ml/10g body weight, i.p. Then, females were decapitated, and brains were rapidly removed from the skull and placed in a cold plaque. There, the HPC and mPFC were identified and dissected. Brain areas were immediately stored at -80 °C until the western blot assay was carried out to evaluate the protein levels of BDNF and phosphorylated tropomyosin receptor kinase B (p-Trkb). Samples were homogenised in a cold lysis buffer, supplemented with protease inhibitor (Complete ULTRA Tablets Mini EASYpack, Roche, Mannheim, Germany) and phosphatase inhibitor (PhosSTOP EASYpack, Roche, Mannheim, Germany). Equal amount of protein (30 μg) for each sample was mixed with loading buffer (153 mM TRIS pH = 6.8, 7.5% SDS, 40% glycerol, 5 mM EDTA, 12.5% 2-β-mercaptoethanol and 0.025% bromophenol blue) and loaded onto 10% polyacrylamide gels, and transferred to PVDF sheets (Immobilion-P, MERCK, Burlington, USA). Membranes were blocked with bovine serum albumin 5% for 30 min at room temperature and then, following an overnight incubation at 4 °C with primary antibodies (See Table 1), membranes were incubated for 2 h at room temperature with their respective secondary fluorescent antibodies (See Table 2). Images were acquired on an Amersham Typhoon NIR Plus laser scanner and quantified using Image Studio Lite software v5.2 (LICOR, USA).

**Table 1.** List of primary antibodies. IHC immunohistochemistry; WB, Western blot.

| Antibody | Description | Assay | Host | Dilution | Company | Item number |
| --- | --- | --- | --- | --- | --- | --- |
| BDNF | Brain-derived neurotrophic factor | WB | Mouse | 1:500 | Abcam | ab205067 |
| pTRKB | Phosphorylated tropomyosin receptor kinase B | WB | Rabbit | 1:500 | Thermofischer | PA5-118577 |
| Tubulin | Tubulin | WB | Mouse | 1:500 | BD Biosciences | 556321 |
| BrdU | 5-Bromo-2'deoxyuridine | IHC | Rat | 1:300 | Abcam | ab6326 |
| NeuN | Neuronal Nuclear antigen | IHC | Mouse | 1:1000 | Merck Millipore | MAB377 |
| VGLUT1 | Vesicular glutamate transporter type 1 | IHC | Guinea pig | 1:1000 | Merck Sigma | AB5905 |
| VGAT | Vesicular GABA transporter | IHC | Rabbit | 1:1000 | Synaptic Systems | 131008 |

**Table 2.**
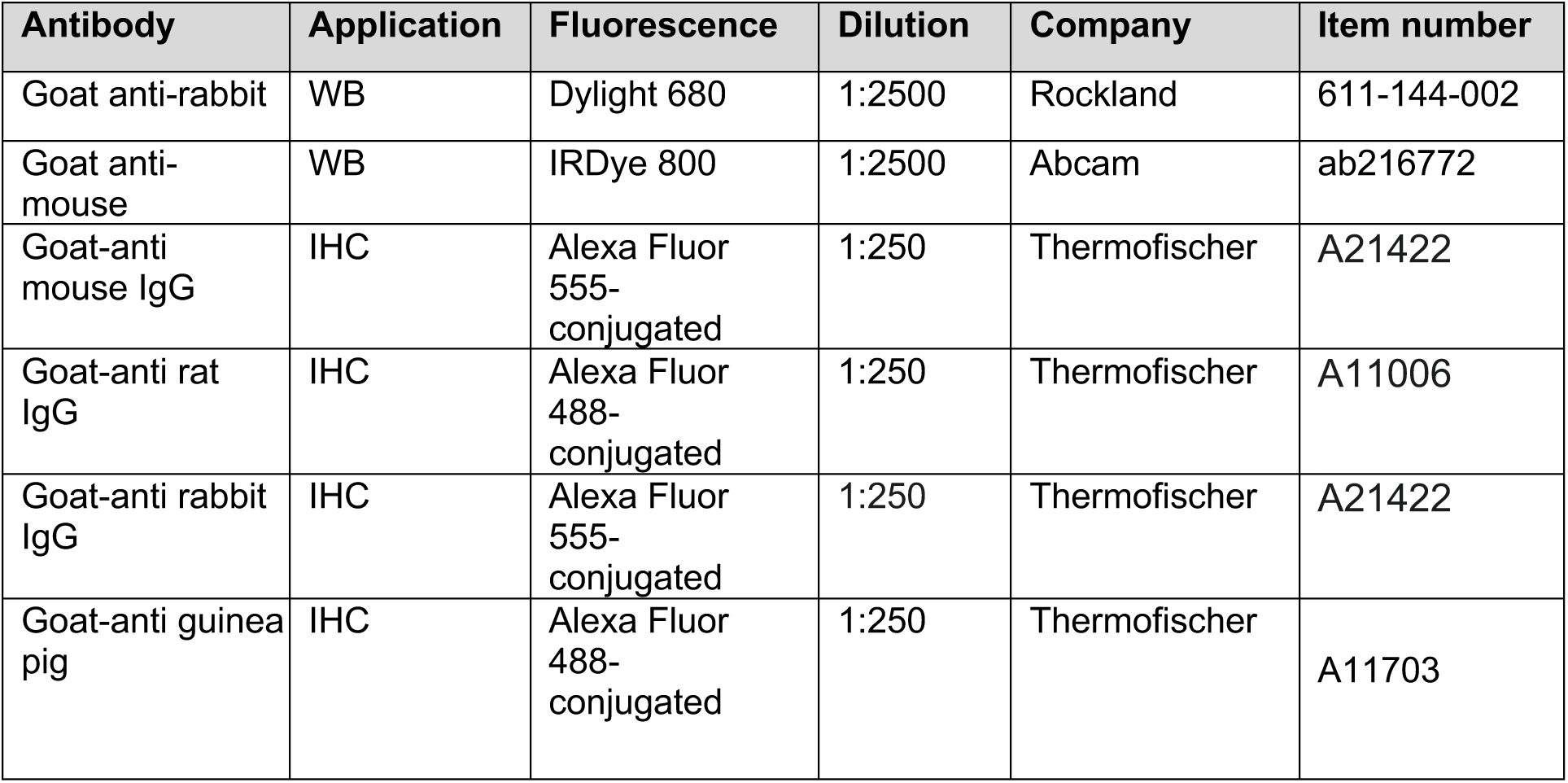
List of secondary antibodies. IHC immunohistochemistry; WB, Western blot.

| Antibody | Application | Fluorescence | Dilution | Company | Item number |
| --- | --- | --- | --- | --- | --- |
| Goat anti-rabbit | WB | Dylight 680 | 1:2500 | Rockland | 611-144-002 |
| Goat anti-mouse | WB | IRDye 800 | 1:2500 | Abcam | ab216772 |
| Goat-anti mouse IgG | IHC | Alexa Fluor 555-conjugated | 1:250 | Thermofischer | A21422 |
| Goat-anti rat IgG | IHC | Alexa Fluor 488-conjugated | 1:250 | Thermofischer | A11006 |
| Goat-anti rabbit IgG | IHC | Alexa Fluor 555-conjugated | 1:250 | Thermofischer | A21422 |
| Goat-anti guinea pig | IHC | Alexa Fluor 488-conjugated | 1:250 | Thermofischer | A11703 |

### 2.8. Determination of endogenous allopregnanolone in plasma

Blood samples from SN (n=8) and MSEW (n=8) dams were obtained immediately after decapitation and collected into EDTA collection tubes (Experiment 1, **Fig. 1A**). Tubes were rapidly centrifuged at 16.400 x *g* for 45 minutes at 4°C, until plasma was separated from blood cells. Supernatant (500 μl) was collected and immediately stored in aliquots at - 80°C until used. Samples selected for ELISA were processed according to the protocol supplied by the ELISA kit manufacturer for allopregnanolone determination (DetectXs Allopregnanolone Enzyme; Immunoassay Kit K044-H1 Arbor Assays, United States). Optical density was read at 450 nm in a Synergy HTX Multi-Mode automatic spectrophotometer. Concentrations were calculated using their standard curves.

### 2.9. Perfusion, fixation and sectioning

One day after BrdU administrations, dams were deeply anaesthetized using a dolethal overdose (i.p. injection of 120 mg/kg of body weight of pentobarbital-based solution). Then, animals were euthanized by transcardiac perfusion of phosphate saline solution 0.1 M (PBS, pH 7.4) using a peristaltic pump (5.5 ml/min for 2 min) followed by 4% paraformaldehyde in 0.1 M phosphate buffer pH 7.4 (same flux for 5 min). Brains were carefully removed from the skull and immediately post-fixed in the same fixative solution overnight at 4 °C. Then, brains were placed into cryoprotectant solution (30% sucrose solution in 0.1 M PBS, pH 7.6, 4 °C) until they sank. Finally, we obtained 30-μm-thick coronal sections using a cryostat (Leica CM3050 S, Wetzlar, Germany). Free-floating sections were collected in five parallel sets.

### 2.10. Immunohistochemistry

Two out of five parallel series obtained from SN and MSEW dams were purposed for immunofluorescence: i) simultaneous of Neuronal Nuclear antigen (NeuN) and BrdU immunostaining in hippocampal sections for cellular proliferation detection, ii) and both vesicular GABA transporter (VGAT, a marker of inhibitory synapses) and vesicular glutamatergic type 1 transporter (VGLUT1, a marker of excitatory synapses) in IL sections adapted by ^34^.

Free-floating sections were washed three times with 0.05 M TRIS buffered saline pH 7.6 (TBS). In brief, sections were: (i) pre-incubated in 4% normal goat serum in 0.05 M TRIS buffered saline pH 7.6 (TBS) with 0.2% Triton X-100, at RT for 1 h, for blocking nonspecific labelling; (ii) incubated in primary antibodies, rat anti-BrdU and mouse anti-NeuN (see Table 1) diluted in TBS with 0.2% Triton X-100 with 4% normal goat serum at RT for 24h; or rabbit anti-VGAT, guinea pig anti-VGLUT1 diluted in TBS with 0.2% Triton X-100 with 4% normal goat serum at 4C for 48h; (iii) incubated with fluorescent-labelled secondary antibodies (90 min at RT) diluted in TBS (See Table 2) for 2h at RT. To reveal the cytoarchitecture in IL sections, they were counterstained prior to mounting by bathing them for 5min in Hoechst (a nuclear staining, 1:10000; Invitrogen, Cat.#33258) at room temperature. After each step, sections were washed three times for 10 minutes in TBS (except between steps (i) and (ii). Finally, sections were washed in TB, mounted onto gelatinized slides and cover-slipped with fluorescence mounting medium (FluorSave Reagent, Merck Milipore, Cat#345789, Darmstadt, Germany). For hippocampal cellular proliferation detection, sections were previously treated with HCl 2N during 30 minutes at 37C. Then, HCl was quenched with borate buffer 0.1M at pH 8.5 for 10 min at RT following^35^.

### 2.11. Analysis of BrdU-positive cells as proliferation cellular marker

The number of BrdU-positive cells was identified using a sequential laser scanning confocal microscopy (Leica TCS SP5 upright). Double scans for NeuN / BrdU-immunostaining, using a 20 × objective, were used to identify Alexa Fluor 488 or Alexa Fluor 555. Sections with 4 μm Z distance of separation were taken from the regions of interest (Bregma -1.46 to -2.30 mm). Photographs were taken in both hemispheres. To minimise the channel spillover, the images were sequentially acquired and saved as LIFF files. To minimise cellular proliferation in the HPC, BrdU-positive cells were manually quantified as total number of labelled body cells in each stack-images per hemisphere using the *cell counter* plugin of Fiji-ImageJ software (Experiment 2, **Fig. 2A**). No manipulations of individual image elements were performed.

### 2.12. Analysis of VGLUT- and VGAT-positive puncta and calculation of E/I ratio

From each immunostaining, slices from the same rostral-caudal level (Bregma 1.70-1.34mm) were examined under a sequential laser scanning confocal microscopy (Leica TCS SP5 upright). Photographs were taken in both hemispheres. Triple (VGAT- / VGLUT1- / Hoechst-immunostaining, using a 63 × oil objective) scans were made to identify Hoechst, Alexa Fluor 488 or Alexa Fluor 555. Sections with 3 μm Z distance of separation were taken from the regions of interest. To minimise channel spillover, images were sequentially acquired and saved as LIFF files. Stacks obtained were further processed with Fiji-ImageJ software. Photographs were taken in both hemispheres. For the analysis of the VGAT- and VGLUT1-positive puncta, the middle 2 optical sections were chosen based on a centre-out focusing strategy as the most high-quality images collected across all cases (2 optical sections/ z-stack, 2 z-stacks/structure).

Then, we perform a 5-step protocol adapting ^36^: i) background substraction from RGB colour images using the sliding paraboloid option, changing the rolling ball (radius set to 50 pixels) to parabolic; ii) enhancement of local contrast by running Contrast Limited Adaptive Histogram Equalization plugin (CLAHE), using block size = 19, histogram bins = 256, maximum slope = 6 as options; iii) application of mathematical exponential (EXP) to further minimise background; iv) execution of the *threshold* command to binarize the image, adjusting it according to an analysis of the histogram of the current image using the 15% of the mean; v) run of the *analyse particles* command, setting size particles= 0.2-infinity. The Excitatory/Inhibitory ratio (E/I) in the neuropil was defined as the number of excitatory puncta divided by the number of inhibitory puncta.

## 3. Statistical analysis

Data were analysed using the software IBM SPSS Statistics 23.0. We first checked the data for normality (Kolmogorov-Smirnov’s test) and homoscedasticity (Levene’s test). We evaluated the nest area, distance to centroid, basal TST, basal splash test, allopregnanolone plasma levels and BDNF/pTrkb protein levels using an unpaired t-Student’s test. We performed two-way ANOVAs for the analysis of the TST, splash test and the immunohistochemical results after treatment injections (with *rearing* and *treatment* as variables in all cases). When applicable, pairwise comparisons were analysed with Bonferroni’s correction. Then, we evaluated differences between groups using the non-parametric Log-rank test (latency to retrieve the first pup in pup retrieval test), Mann-Whitney *U* test for the number of retrieved pups and Fischer’s test for the proportion of mothers that retrieved the first pups within the first 600 s, pups retrieved and the number of females that retrieved all pups to the nest. Statistical differences were found when *p* < 0.05.

## 4. Results

### 4.1. Characterization of depressive-like phenotype in MSEW postpartum mice

As a core symptom of PPD, we analysed the maternal behaviour of postpartum MSEW and SN females. In the pup retrieval experiment, results of the Log-rank test revealed that all SN lactating females retrieved the first pup within the 600 s, whereas only 50% of MSEW females did retrieve the first pup (**Fig. 1B**; χ_2_ = 5.64, *p* < 0.05). When we analysed the number of retrieved pups, we observed highly significant differences between SN and MSEW groups (**Fig. 1D**; Mann-Whitney, *p* < 0.001), since most SN mothers retrieved all their pups, whereas only 2 of MSEW females did (**Fig. 1C**; Log-Rang test, χ_2_ = 5.64, *p* < 0.01). Consistent with deficits in pup retrieval behaviour, MSEW mice showed larger distances to nest centroid (**Fig. 1I**; t_19_ = 2.41, *p* < 0.05) and bigger nest areas (**Fig. 1H**; t_19_= 4.15, p < 0.001) than SN (**Fig. 1E**), suggesting that pups are scattered randomly throughout the cage in MSEW litters as **Fig. 1F** and **G** shows. Finally, MSEW females displayed increased digging behaviour as a measure of stress-coping behaviour in comparison with SN (**Fig. 1J**; t_19_ = 5.19, *p* < 0.001).

On PW 2, MSEW and SN mothers underwent the TST to evaluate despair-like behaviour as a measure of depressive phenotype. Results indicate that MSEW dams displayed longer immobility time compared to SN (**Fig. 1K**; t_17_ = 2.45, *p* < 0.05). On PW 5, the splash test was performed to evaluate self-care and hedonic behaviour ^33^, to further characterise the depressive-related phenotype of MSEW dams. Females exposed to MSEW exhibited a decreased self-grooming behaviour when compared to SN mothers (**Fig. 1L**; t_18_ = 3.15, *p* < 0.01).

### 4.2. Allopregnanolone plasma levels in postpartum female mice

Since low circulating levels of allopregnanolone are associated with PPD ^17^, we investigated whether MSEW could reduce the plasma content of allopregnanolone. On PW 2, we found a general tendency towards decreased allopregnanolone plasma levels in MSEW postpartum females when compared to their SN counterparts (**Fig.1M**; t_14_ = 2.04, *p* = 0.06).

### 4.3. MSEW induces a reduction of p-TRKB marker in PFC of postpartum females

Brain tissue was extracted to determine whether MSEW could induce changes in the activation of BDNF-Trkb pathway. In the PFC, t-Student’s test analysis showed that pTRKB protein levels were reduced in MSEW postpartum females (t_15_= 2.15, *p* < 0.05; **Fig.1N**) but no significant differences were observed in BDNF expression (**Fig.1O**) between MSEW and SN groups. By contrast, when we analysed the expression of these proteins in the HPC, statistical differences were not found (**Fig.1P** and **Q**).

### 4.4. Ketamine and allopregnanolone reduces postpartum depressive-like behaviour in MSEW dams

In order to address whether ketamine and allopregnanolone reduce despair-like behaviour, SN and MSEW dams were subjected to TST (PW 2 **Fig. 2A**) after treatments. The analysis of TST results using a two-way ANOVA revealed a *treatment* effect (F_2,60_ = 7.82, *p* < 0.01) and *rearing* × *treatment* interaction (F_2,60_ *=* 10.16, *p* < 0.001; **Fig. 2B**). The *treatment* effect indicated that both ketamine (*p* < 0.01) and allopregnanolone (*p* < 0.001) diminished the immobility time in comparison with vehicle-treated postpartum females. After Bonferroni’s correction, results showed that vehicle-treated MSEW mice presented higher immobility time (*p* < 0.001) compared to the vehicle-treated control animals. Both ketamine (*p* < 0.001) and allopregnanolone (*p* < 0.001) decreased the immobility time in MSEW postpartum mice.

### 4.5. Only ketamine restores self-grooming behaviour in MSEW dams

As many clinicians recognised, anhedonia constitutes a major symptom of depression and, for this reason, it is an important measure of depressive-like behaviour in animal models. Female mice underwent the splash test to study anhedonia after ketamine and allopregnanolone treatments on PW 5 (**Fig. 2C**). Two-way ANOVA analysis indicated *rearing* × *treatment* (F_2,60_ = 7.65; *p* < 0.01) effect. The *post hoc* Bonferroni’s analysis showed vehicle-treated MSEW dams displayed less self-grooming behaviour when compared to vehicle-treated SN mice (*p* < 0.001), whereas ketamine-treated MSEW females restored self-care behavioural deficits shown in vehicle-treated MSEW females (*p* < 0.01). By contrast, allopregnanolone was not able to ameliorate said impairments, so ketamine-treated MSEW females showed higher self-grooming behaviour (*p* < 0.05). These results, indicate that both treatments exert antidepressant effects on MSEW female mice, however, only ketamine was effective in reducing anhedonia present in MSEW females.

### 4.6. Postpartum depression decreases neural hippocampal proliferation in females: role of ketamine and allopregnanolone

Following the behavioural assessment, we investigated neuronal proliferation in the dentate gyrus of HPC (**Fig. 3A**). The analysis of the number of BrdU-positive cells using a two-way ANOVA showed *rearing* (F_1,26_ = 6.29; *p* < 0.05) and *treatment* (F_2,26_ = 16.86; *p*< 0.001) effects (**Fig. 3B**). The *rearing* effect suggests that MSEW female mice (**Fig. 3D, F** and **H**) had lower hippocampal proliferation than the SN group (**Fig. 3C, E** and **G)**. Regarding *treatment* effect, ketamine-treated animals (**Fig. 3E** and **F**) showed an increased cellular proliferation when compared to vehicle-treated groups (*p* < 0.05; **Fig. 3C** and **D**). Moreover, allopregnanolone treatment induce a higher cellular proliferation in female postpartum mice compared to vehicle- (*p* < 0.001; **Fig. 3C** and **D**) and ketamine-treated animals (*p* < 0.05; **Fig. 3E** and **F**).

**Fig. 3.**
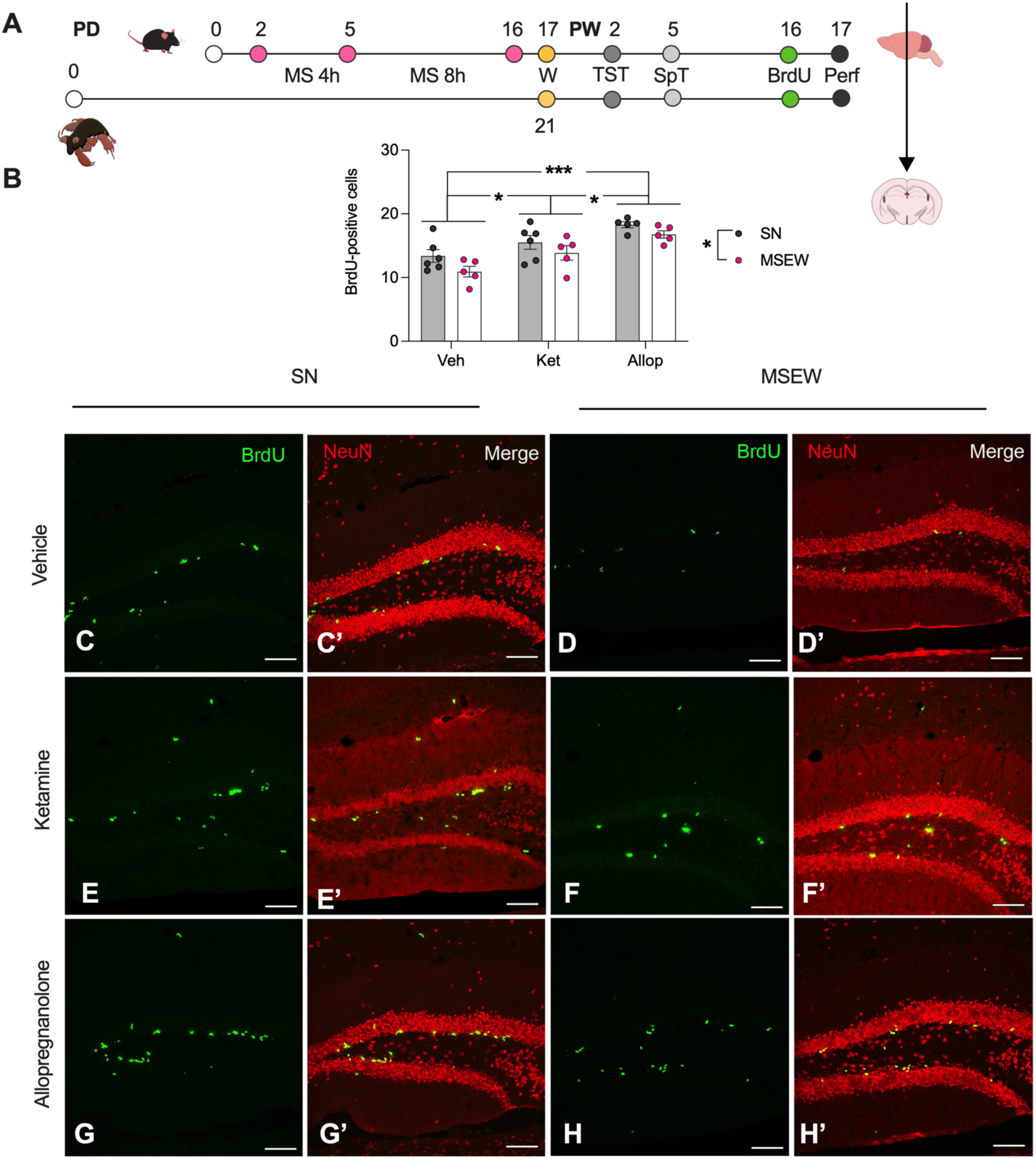
Increased hippocampal cellular proliferation induced by ketamine and allopregnanolone. (A) Timeline of the experiments. Grey (SN) and pink (MSEW) dots represent (B) the average of BrdU-positive cells per hemisphere after vehicle, ketamine or allopregnanolone treatments (n = 5-6 / group). Representative of single stack of BrdU (green cells) and NeuN-BrdU merge of vehicle- (images of SN female are C and C’; images of MSEW female are D and D’), ketamine- (images of SN female are E and E’; images of MSEW group are F and F’) and allopregnanolone-treated animals (images of SN female are G and G’; images of MSEW female are H and H’). All data are represented as the mean ± SEM. (Two-way ANOVA _*_p < 0.05, ^***^p < 0.001). Scale bar 100 μm. Allop, allopregnanolone; BrdU, 5-bromo-2’Deoxyuridine; Ket, ketamine; MS, maternal separation; MSEW, maternal separation with early weaning; Perf, perfusion; PD, postpartum day; PW, postweaning day; SN, standard nest; SpT, splash test; TST, tail suspension test; W, day of weaning day.

### 4.7. MSEW decreases VGAT puncta without altering E/I ratio

We therefore sought to understand how excitatory and inhibitory synapses could be altered by MSEW, as well as whether ketamine and allopregnanolone could restore those deficits. To that end, we evaluated the number of puncta VGAT and VGLUT1 in the IL (**Fig. 4A**). Two-way ANOVA analysis for the number of VGLUT1-positive puncta revealed significant differences due to *treatment* (F_2,29_ = 11.78; *p* < 0.001) and a trend in *rearing* effect (F_2,29_ = 3.55; *p* = 0.07), indicating a decreased expression of VGLUT1 puncta in MSEW females (**Fig. 4F, H** and **J**). When we analysed the effect of *treatment* (**Fig. 4B**), we found that allopregnanolone increased the number of VGLUT1-positive puncta in the IL (**Fig. 4I** and **J**) in comparison with vehicle (*p* < 0.01; **Fig. 4E** and **F**) and ketamine (*p* < 0.001; **Fig. 4G** and **H**) treatments. Regarding VGAT-immunoreactive puncta, we found significant differences due to *treatment* (F_2,29_ = 14.68; *p* < 0.001) and *rearing* factors (F_2,29_= 9.35; *p* < 0.01; **Fig. 4C**), indicating that MSEW reduced the number of inhibitory VGAT puncta in the IL (**Fig. 4F’, H’** and **J’**). As for the *treatment* effect, *post hoc* Bonferroni’s analysis showed that allopregnanolone (**Fig. 4I’** and **J’**) increased the number of VGAT-positive puncta when compared to vehicle (*p* < 0.001; **Fig. 4E’** and **F’**) and ketamine (*p* < 0.001; **Fig. 4G’** and **H’**) treatments. However, statistical differences were not found when we analysed the E/I ratio (**Fig. 4D**).

**Fig. 4.**
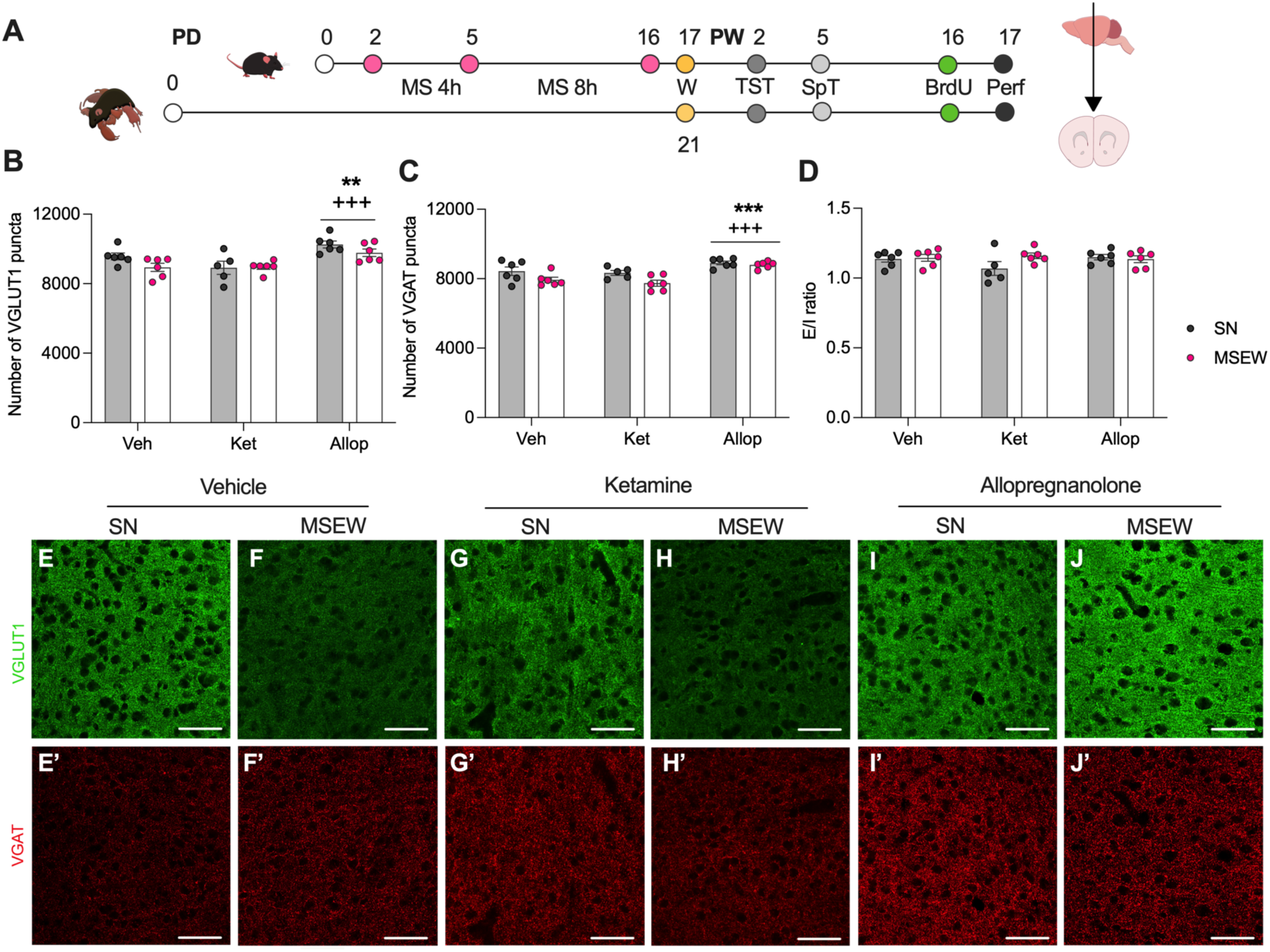
MSEW decreases the number of VGAT puncta in the infralimbic cortex. (A) Timeline of the experiments. Grey (SN) and white (MSEW) bars represent the average number of (B) VGLUT1-, (C) VGAT-positive puncta in IL and (D) excitatory/inhibitory ratio (n = 5-6 / group) after vehicle, ketamine or allopregnanolone treatments measured in two different confocal z-planes in both hemispheres of each individual animal. Representative single confocal planes of puncta expressing excitatory (green, VGLUT1, E-J) and inhibitory (red, VGAT, E’-J’) synaptic markers in IL. All data are represented as the mean ± SEM. (Two-way ANOVA ^*^p < 0.05, ^***^p < 0.001). Scale bar 50 μm. Allop, allopregnanolone; BrdU, 5-bromo-2’Deoxyuridine; Ket, ketamine; MS, maternal separation; MSEW, maternal separation with early weaning; Perf, perfusion; PD, postpartum day; PW, postweaning day; SN, standard nest; SpT, splash test; TST, tail suspension test; VGAT, vesicular GABA transporter; VGLUT1, vesicular glutamate transporter type 1W, day of weaning day.

## 5. Discussion

The present study constitutes a detailed characterisation of a new model of PPD-like phenotype induced by MSEW in female C57BL/6 mice, which confers a complete recapitulation of the core symptoms of PPD. In this sense, postpartum females subjected to MSEW display an increased despair-like behaviour and decreased anhedonia by means of the TST and splash test, respectively. Several studies have extensively reported depressive- and anhedonia-like behaviours in postpartum rats that underwent long-term maternal separation (for review see ^37^), but this is the first time this is described in mice. Moreover, MSEW dams present a maternal neglect via pup retrieval test. In fact, 50% of MSEW dams do not retrieve all pups to the nest and they display repetitive disruptive digging behaviour.

We have previously reported that MSEW protocol causes persistent detrimental consequences in offspring development, inducing behavioural and neurobiological alterations ^27,28,38^, that also might occur in children whose mother who has experienced PPD ^39,40^. Altogether, these findings highlight the importance of a mother-infant relationship for a healthy development of both parts.

Among the multiple factors potentially involved in the development of PPD, neurosteroids that acts as positive allosteric modulators ^14,41^ and neurotrophins ^42,43^, such as allopregnanolone and BDNF-Trkb intracellular pathway, have been proposed as key biomarkers for PPD. In line with women who have experienced PPD ^17,44^ and maternity ‘blues’ ^45^, MSEW dams show a long-term reduction of allopregnanolone plasma content. This could be due to the stress induced by MSEW protocol given that repeated exposure to stress alters the hypothalamo-pituitary-adrenal (HPA) axis and dampens allopregnanolone release ^46^. Nevertheless, other studies have not found differences in circulating levels of allopregnanolone ^8^, but probably due to the time point of blood sample extraction.

As per BDNF, we wanted to explore its expression levels in specific brain regions of MSEW model, since lower levels in serum of late pregnant and postpartum women correlate with higher depressive symptoms ^42,43^. In agreement with other rodent animals of postpartum depression ^47,48^, we did not observe changes in the expression of BDNF in MSEW in either PFC or HPC. However, this is controversial because decreased hippocampal BDNF immunoreactivity has been dpcumented in other models of PPD, like gestational stressed rats ^49^. Different time points sample extraction or analytical techniques might account for the reported differences. Nevertheless, we did find decreased protein levels of phosphorylated-Trkb only in PFC of MSEW mice, indicating downregulation of the Trkb signalling pathway. Accordingly, decreased Trkb activation pathway within PFC has been correlated with depressive-like symptoms in different stress-induced models of depression, probably caused by the high levels of glucocorticoids due to stress ^50^.

Additionally, in our study we compared the expression of VGLUT1 and VGAT in IL between MSEW and SN females. These transporters mediate the synaptic release of glutamate and GABA, respectively ^51,52^, therefore the efficacy of synaptic neurotransmission ^51^. MSEW dams show decreased VGLUT1 and VGAT positive puncta in IL, as previously reported in other animal models of depression ^53,54^. Deficits of GABAergic neurotransmission, as well as abnormal expression of GABA_A_ receptors have been described as part of the pathophysiology of PPD ^16,18^. In fact, a GABA_A_ knock out postpartum females display maternal neglect, cannibalism and higher depressive-like symptoms^18^. In this respect, chronic stress leads to a maladaptive balance of GABAergic and glutamatergic signalling, but the excitatory and inhibitory ratio seems to remain somehow balanced after chronic stress due to allostatic regulation. Nevertheless, those synapses are thought to remain impaired which would explain the reduced neural connectivity observed in PPD ^16^. Our results support this hypothesis because, although both VGLUT1 and VGAT were reduced, the E/I was not different between SN and MSEW dams.

Likewise, we have analysed adult hippocampal neurogenesis counting BrdU-positive cells, a marker of proliferative neurons. MSEW reduced dorsal hippocampal proliferation in postpartum females. Compelling evidence from preclinical studies has suggested a functional link between neurogenesis, stress-related behaviours and depression ^12,55,56^. Targeting adult hippocampal neurogenesis does not directly exert antidepressant effects, but it might counteract the disruptions at behavioural and circuitry levels induced by chronic stress through modulation of the HPA axis ^55^. This might indicate that adult hippocampal neurogenesis participates in the physiological function of the endocrine system and the behavioural component of the stress response. Since stress could be considered as a major trigger for PPD, we hypothesise that MSEW-induced stress promotes adult hippocampal neurogenesis impairments underlying a PPD-like phenotype together with a poorer stress responsivity. In fact, lower endogenous allopregnanolone serum levels, as we found in MSEW postpartum females, have been associated to dysregulated HPA axis and altered hippocampal neurogenesis ^57^.

Currently, there is no effective treatment to achieve the complete symptomatology remission of all PPD patients. Even though exogenous allopregnanolone (Brexanolone, Zulresso) has been recently approved as a unique treatment for PPD by the Food Drug Administration ^19,20^ and ketamine has been used as a prophylactic treatment after caesarean sections ^22,58^, the exact mechanism of action of both drugs remains barely speculative. Here we demonstrated that low doses of ketamine reverse the depressive-and anhedonia-like behaviour induced by MSEW, as previously shown in other animal models of depression ^59,60^. However, we did not find an effect of ketamine on glutamatergic and GABAergic signalling markers within IL using immunohistochemical techniques. Recently, it has been proposed that sub-anesthetic doses of ketamine may exert theis antidepressant effects by enhancing the activity and E/I input of pyramidal neurons as well as decreasing the number of interneurons located in the prefrontal cortex ^61^. Nonetheless, those analyses were carried out using electrophysiological approaches, hence, perhaps we are unable to detect this fine-tune modulation due to the sensitivity of the technique used. By contrast, exogenous allopregnanolone exerts antidepressant effect as previously documented ^30,62^, without affecting anhedonic-like behaviour. Nevertheless, allopregnanolone increases both VGLUT1- and VGAT-positive puncta within IL in comparison with ketamine- and vehicle-treated animals. Given that both glutamatergic and GABAergic synaptic markers were enhanced, E/I ratio remained unaltered, suggesting that allopregnanolone treatment might bring the system towards the homeostatic conditions in the MSEW dams. These findings support previous results in the literature whose mechanism of action is by increasing GABAergic neurotransmission ^16,63^ and, indirectly modulating glutamatergic release in IL, as it also occurs in other brain areas^64^.

Finally, our results indicate that ketamine and allopregnanolone are pro-neurogenic treatments, because both drugs enhanced the number of BrdU-positive cells in dorsal hippocampus of all the females, in accordance with previous studies ^12,57,65^. Increased neural proliferation in the hippocampus might be indirectly participating in the amelioration of depressive-like behaviours by decreasing the hyperactivity of the HPA axis, as previously documented in other stress-induced mouse models of depression ^55^.

## CONCLUSION

MSEW protocol represents a reliable behavioural model to investigate certain aspects of PPD. Postpartum female mice that undergo MSEW protocol exhibit depressive-like phenotype after weaning, such as despair- and anhedonic-like behaviours assessed by the TST and splash test respectively. Moreover, these dams show neglecting maternal care, a core symptom of PPD.

Even though exogenous allopregnanolone administration is the current major treatment for PPD, our data indicate that ketamine might proof to be even more effective than allopregnanolone in our mouse model, because ketamine is able to reduce both despair-like and anhedonic-like behaviours, whereas allopregnanolone only impacts the despair-like domain. Both drugs induce pro-neurogenic effects within the HPC, supporting the hypothesis of antidepressant drugs acting through hippocampal neurogenesis. In addition, allopregnanolone increases VGAT and VGLUT markers in the IL, while such low dose of ketamine does not affect their expression. Overall, our findings have shed some light on the promising effects of ketamine on PPD-like subjects and, therefore, we claim the need of further studies exploring the effectiveness of low doses of ketamine on PPD patients.

## 6. Authors contribution

A.M-S. and O.V. were responsible for the study concept and design. A.G-B, X.P-R., I.F-G., I.G-L. and A.M-S. carried out the experimental studies. A.G-B., A.M-S. and O.V. participated in the interpretation of findings. A.G-B., A.M-S and O-V. drafted the manuscript. All authors critically reviewed the content and approved the final version for publication.

## 8. Acknowledgements

The authors are indebted to Javier Valle-García (Proteomics and Protein Chemistry Group; Universitat Pompeu Fabra), Marc González-Colell (Synthetic Biology for Medical Application, Universitat Pompeu Fabra), Elia Lanàquera-Ivars and Irene Rodríguez-Navarro for their invaluable assistance during the present study.

## 9. Abbreviations

BDNF: Brain-derived neurotrophic factor
BrdU: 5-Bromo-2’deoxyuridine
E/I: Excitatory/Inhibitory ratio
GABA: γ -aminobutyric acid
HPA: Hypothalamus-pituirary-adrenal
IL: Infralimbic cortex
MSEW: Maternal separation with early weaning
NeuN: Neuronal Nuclear antigen
p-Trkb: Phosphorylated Tropomyosin receptor kinase B
PD: Postpartum day
PPD: Postpartum Depression
PW: Postweaning day
SN: Standard Nest
TST: Tail suspension test
VGAT: Vesicular GABA transporter
VGLUT1: Vesicular glutamate transporter type 1

